# Transmission of allosteric response within the homotrimer of SARS-CoV-2 spike upon recognition of ACE2 receptor by the receptor-binding domain

**DOI:** 10.1101/2020.09.06.284901

**Authors:** Sayan Bhattacharjee, Rajanya Bhattacharyya, Jayati Sengupta

## Abstract

The pathogenesis of novel SARS-CoV-2 virus initiates through recognition of ACE2 receptor (Angiotensin-converting enzyme 2) of the host cells by the receptor-binding domain (RBD) located at spikes of the virus. Following receptor-recognition, proteolytic cleavage between S1 and S2 subunits of the spike protein occurs with subsequent release of fusion peptide. Here, we report our study on allosteric communication within RBD that propagates the signal from ACE2-binding site towards allosteric site for the post-binding activation of proteolytic cleavage. Using MD simulations, we have demonstrated allosteric crosstalk within RBD in apo- and receptor-bound states where dynamic correlated motions and electrostatic energy perturbations contribute. While allostery, based on correlated motions, dominates inherent distal communication in apo-RBD, electrostatic energy perturbations determine favorable crosstalk within RBD upon binding to ACE2. Notably, allosteric path is constituted with evolutionarily conserved residues pointing towards their biological relevance. As revealed from recent structures, in the trimeric arrangement of spike, RBD of one copy interacts with S2 domain of another copy. Interestingly, the allosteric site identified is in direct contact (H-bonded) with a region in RBD that corresponds to the interacting region of RBD of one copy with S2 of another copy in trimeric constitution. Apparently, inter-monomer allosteric communication orchestrates concerted action of the trimer. Based on our results, we propose the allosteric loop of RBD as a potential drug target.

---

The recent outbreak of a novel coronavirus, the SARS-CoV-2 (severe acute respiratory syndrome coronavirus 2), a close relative of the previously-known SARS-CoVs, has become an ongoing global public-health threat. Following its emergence, it has created an pandemic situation within no time owing to its alarming high rate of transmission^1^.

Cell entry in particular, is an important early step of cross-species transmission. Spike glycoprotein of CoVs plays the role of attachment to the Angiotensin-converting enzyme-2 (ACE2) receptor of the host cells and mediates viral entry. The association with the host-cell receptor is regulated by the receptor-binding domain (RBD) of the spike protein followed by cleavage using a host protease, which releases the spike fusion peptide, and thereby promotes viral entry^2^.

SARS-CoV-2 is a single-stranded RNA virus with four major constituent proteins: spike protein (S), membrane protein (M), envelope protein (E), and nucleo-capsid protein (N)^3^. Lung is the key target organ of the virus and, as the name implies, the infection induces pneumonia accompanied by acute respiratory distress.

Recent high resolution structures of the trans-membrane spike protein of SARS CoV-2 show^4^ intertwined arrangement of three monomers. Each monomer consists of sub-domains S1 and S2. The receptor-binding domain (RBD) that interacts with ACE2 receptor of the host belongs to the S1 subunit, while the S2 subunit anchors on the viral membrane. Interestingly, RBD of one copy directly interacts with the S2 domain of another copy in the homotrimer (Figure 1). However, the receptor binding motif (RBM) in the RBD that interacts with the receptor has not been fully resolved in the recent spike trimer structures. A following crystallographic study^5^ has resolved the interactions of SARS-CoV-2 spike RBD with the ACE2 receptor.

**Figure 1:**
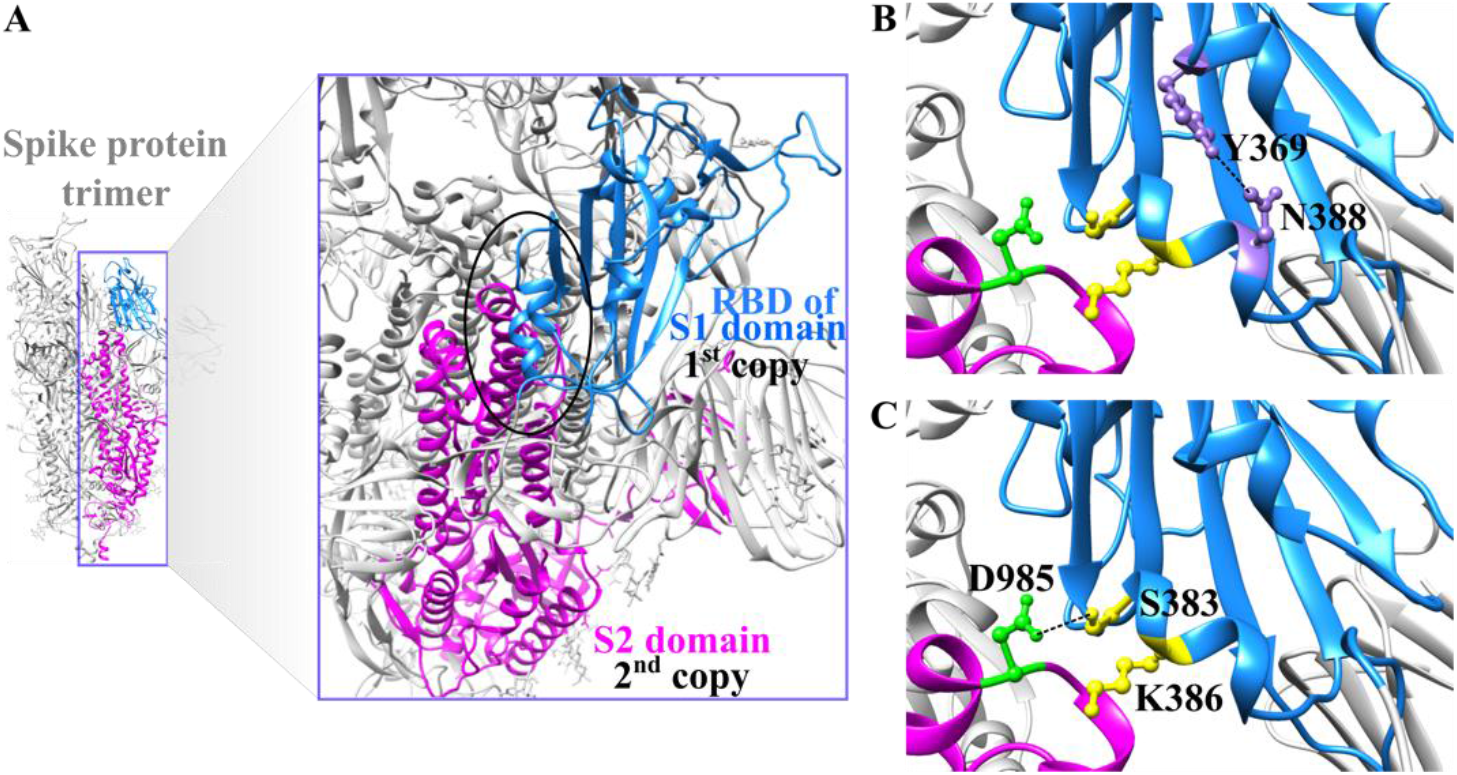
RBD of S1 and S2 domain of Spike protein. (**A**) Shows trimeric Spike protein structure (PDB ID: 6Z43). The RBD of the S1 domain of one monomer (light blue) and the S2 domain of other monomer (magenta) is shown in close up view. The allosteric region of RBD (part of S1 domain of one monomer, marked as 1^st^ copy) and the S2 domain (neighboring copy of monomer, marked as 2^nd^ copy) connection is marked with black circle to show the post-receptor binding favorable allosteric signal transfer may occur in the trimeric Spike protein, as predicted from the electrostatic perturbations in RBD. (**B**) Represents further close up view of the two allosteric loops connected through H-bond. (**C**) Communication between RBD of the 1^st^ copy and S2 domain of the 2^nd^ copy is shown with both H-bond and electrostatic interaction. H-bonds are shown in broken black lines.

Clearly, successful viral infection is reliant on the effective complex formation between the host receptor and the protruded spikes on the viral surface, which makes it an obvious drug target. However, in addition to the contact residues, studies have indicated potential role of non-contact residues and allostery in the spike protein-receptor interactions^6^.

To better understand the etiology of the infection, we have focused on the RBD of the spike protein and its binding to the ACE2 receptor^6a, 7^, information on which may provide important clues for designing drugs to combat the virus. Our study aims to understand the hidden allosteric cross-talks in RBD upon recognition of the ACE2 receptor and their possible role in the activity of the spike protein trimer. The allosteric crosstalk between the ACE2-binding residues of RBD with its distal residues (away from the binding site) was evaluated using MD simulation followed by extensive analyses of both dynamics and energy terms at atomic level. MD simulation has long been implicated as a suitable procedure to explore the allosteric paths of information exchange in between amino acid residues of a protein by elucidating intermediate protein conformations that often occur in pico or nano second time scale which are often difficult to capture with other techniques^8^. The conformational dynamics and change in energetic within RBD before and after binding to the host receptor revealed the hidden communication that shapes the crosstalk pathway leading to recognition and subsequent internalization of the virus.

Our study offers the identified allosteric sites as additional, potential drug targets. Insights gained from our results on electrostatic contributions in viral infection suggest that modification or inhibition of the allosteric communication would block proper functioning of the viral spike protein and consequently hinder viral infection. Therefore, it may be considered as possible intervention pathway for infection prevention.

## Results and Discussion

### Conformational dynamics of free and receptor-bound RBD

To probe deeper into the dynamicity of free and ACE2-bound RBD of the COVID spike protein, Cα atoms based root mean square fluctuations (RMSF) and Order parameter (*S^2^*) considering the N-H vectors^9^ of RBD, were calculated from the corresponding trajectories. The overall bound state RMSF revealed a lower value relative to the free RBD, implicating the higher rigidity of RBD after binding to ACE2 compared to its apo-form. This observation suggests the ACE2-directed stabilization of the RBD in the bound state by selection of particular conformation from the fast time scale (ps-ns) regime.

In this regard, the residues exhibiting remarkably differentiated RMSF values between the two states are clearly regions displaying some motifs having higher plasticity in free RBD (an RMSF cut-off of >0.25 nm was set to define regions of high flexibility): the binding loop (containing residues 474 to 486), and a distal loop (consisting residues 358 to 376). It is apparent that the distal loop can be considered as an allosteric site (away from the ACE2 binding site of RBD). Remarkably, this allosteric loop is in direct contact through H-bond to a region of the RBD that coincides with the region of one monomer that interacts with the S2 domain of another monomeric copy in the spike protein trimer^4a, 10^ (Figure 1).

Significantly, when the dynamicity of the apo-form was compared with respect to the ACE2-bound RBD, the fluctuation of the allosteric loop decreased considerably along with the ACE2 binding loop (Figure 2A-C). This diminution in fluctuation was expected for the binding loop but simultaneous decrement in dynamicity of the distal loop pointed towards allosteric involvement through correlated motion^11^. Interestingly, a slightly enhanced flexibility was observed in another distal loop (containing residues 384 to 390) of RBD (which actually involved in interactions with S2 of another copy (Figure 1)) upon binding to ACE2 (Figure2).

**Figure 2:**
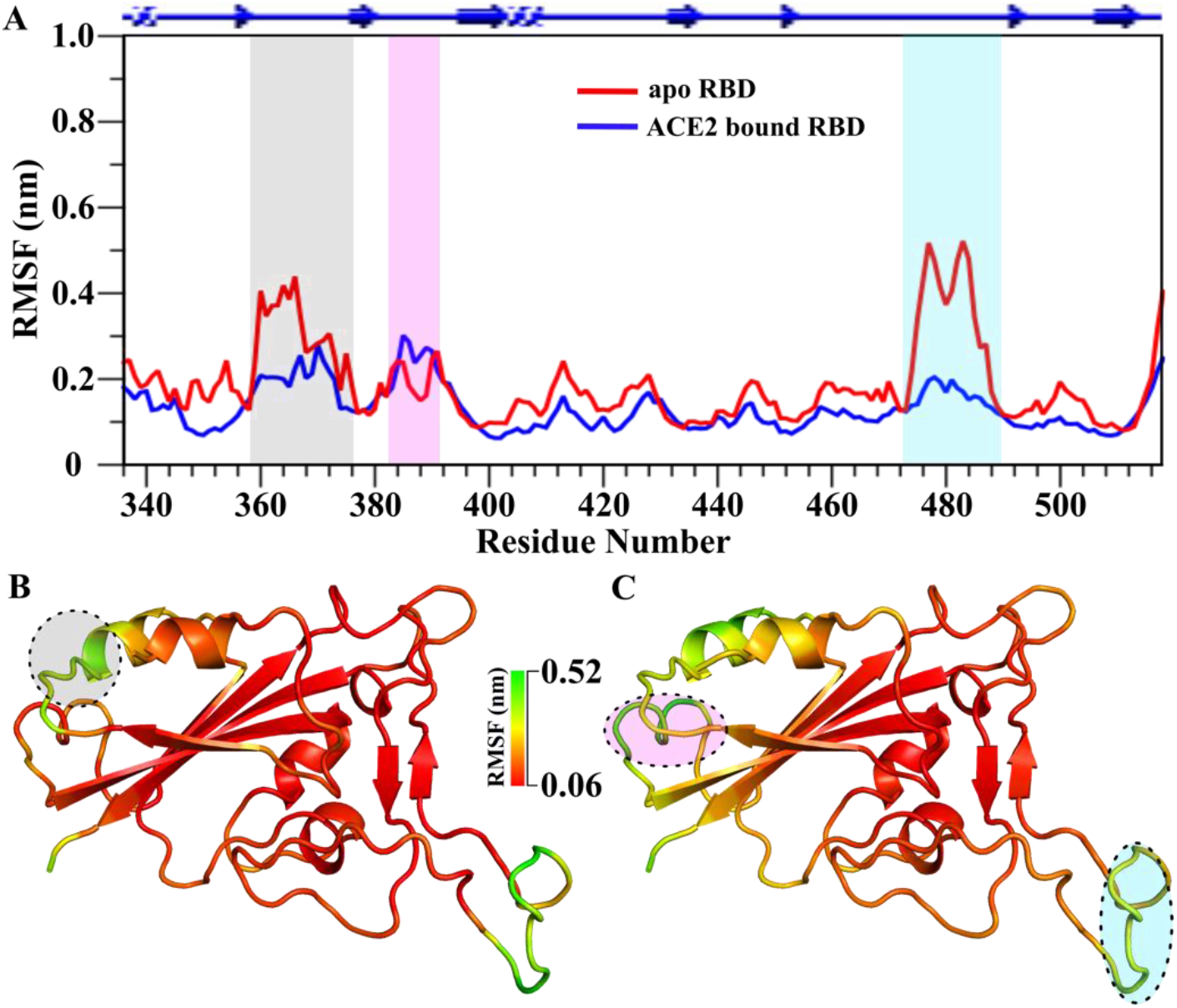
Comparison of Cα atom dynamicity. (**A**) Residue wise Cα atom root mean square fluctuations for both free and ACE2 bound RBD of SARS-Cov-2. Significant changes (above cut-off value of 0.2 nm) were marked with different colored columns. RMSF values were mapped onto the crystal structures (PDB ID: 6M0J) of respective (**B)** free, and (**C**) ACE2-bound RBD. A spectrum bar based color scheme is shown (using PyMol) to point out differential fluctuating regions of both, and are indicated by circles corresponding to the color scheme used in (**A**).

Analysis of order parameters (*S^2^*) based on N-H vectors singularly complied with the conclusions of the RMSF analysis where similar allosteric fluctuations (in both the above mentioned distal loops) were also identified, with an overall decrement in the dynamicity of ACE2-bound RBD with substantial alterations in the flexibility of the binding region (between residues 474 to 486)(Figure S2). The *S^2^* parameter represents flexibility and rigidity of a protein with the values from 0 to 1, where values close to 0 and 1 indicate high flexibility and rigidity respectively. Thus, this observation also suggests the decrement in overall RBD backbone entropy factor (*S^2^* is directly related to conformational entropy^12^) and hence formation of a stable complex.

### Modulation of allosteric communication within RBD upon receptor binding

Allosteric involvement in proteins is one of the most important areas in the field of allosteric drug development and it was proposed that allosteric communications were achieved either by correlated motions of the amino acids or by fundamental non-bonded energy contributions^11, 13.^

A number of noticeable differences in correlation were found when DCCM (a residue-wise correlation matrix-based approach where the correlated and anti-correlated motions are scored between 1 and −1 considering strong: |*C_ij_*| = 1.0–0.7; moderate: |*C_ij_*| = 0.7–0.5; and weak: |*C_ij_*| = 0.3 −0.5) of both apo- and ACE2-bound RBD was compared. Free RBD consisted of several strongly correlated and moderately anti-correlated motions within the amino acids, which were either missing or became weakly correlated in the ACE2-bound RBD (Figure 3A-B). Interestingly, the binding loop (residues from 474 to 486) and the allosteric loop (residues from 358 to 376) showed a moderate anti-correlated motion. In other words, these loops move away from each other in a correlated manner. It may be noted here, both the allosteric loops are in correlation which reestablishes the presence of H-bond between them (Figure 1B) as revealed by crystal structure described above. This kind of anti-correlated motion, which could be attributed to the structural flexibility of apo-RBD, is much weakened in the ACE2-bound RBD owing to its conformational rigidity in bound state.

**Figure 3:**
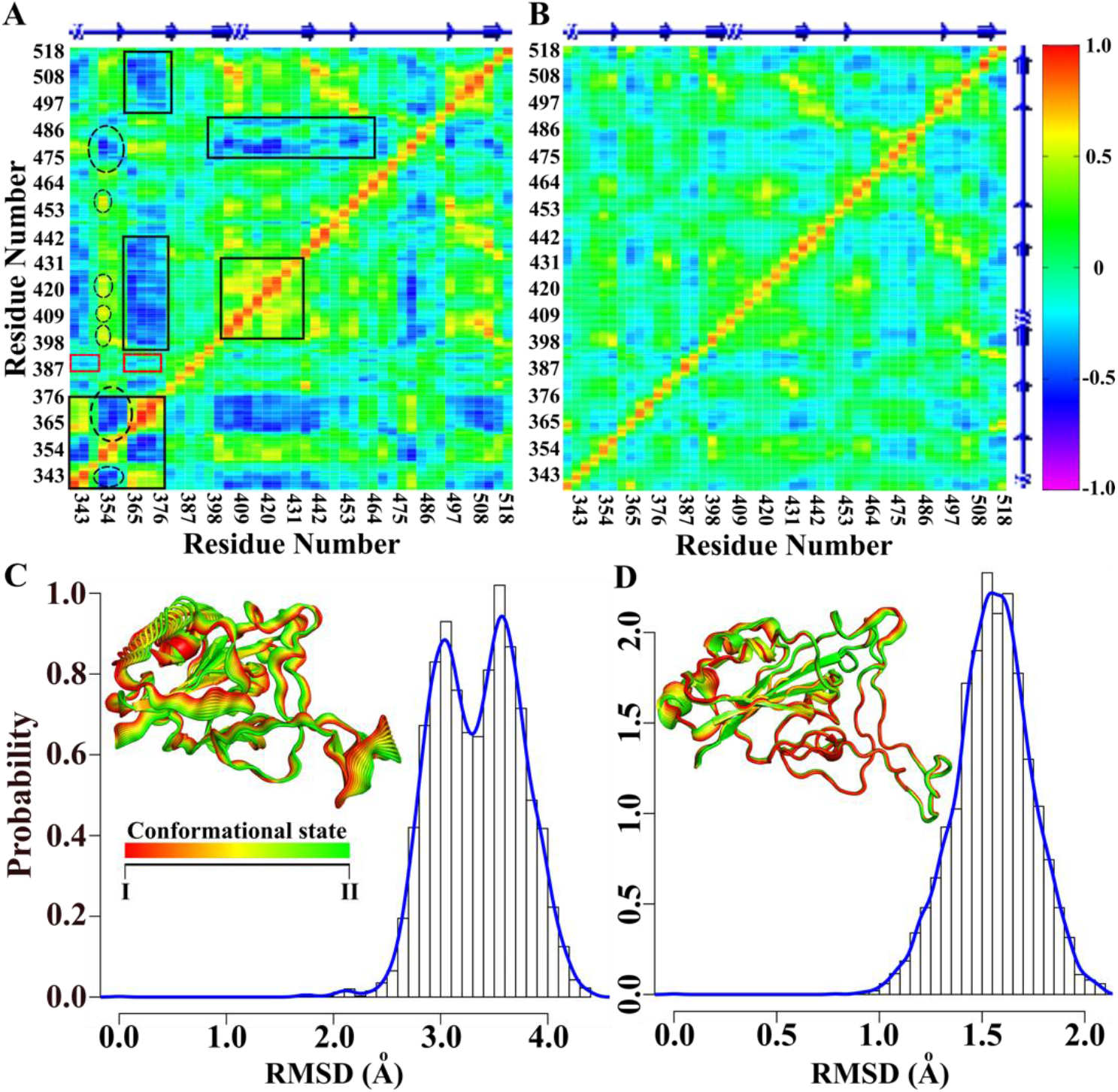
Correlated motions and dynamicity. DCCM of (**A**) apo and (**B**) ACE2 bound RBD are shown. Differential residue wise correlations in both the form of RBD are indicated by boxes. Allosteric communication path was marked with broken circle. Correlation between two allosteric loops is shown in small red box. RMSD probability distribution reflected two populations in apo- (**C**), and single population in ACE2-bound RBD (**D**). The 3D projections of both are shown in insets (calculated from PCA). Apo-RBD dynamicity reflected conformational alterations between two states from ps-ns time scale regime of dynamicity.

For further verification, Root Means Square Deviations (RMSD) distribution and Principal Components Analysis (PCA) of both free and bound RBD were calculated. The RMSD distribution comparative study between the two forms revealed strikingly divergent populations. Two distinct populations were observed for the apo-RBD that were in agreement with PCA results while the bound RBD noticeable displayed a single RMSD population (Figure 3C-D and S3A-B).Both RMSD and PCA results clearly indicated the flexible conformational alterations of apo-RBD between two conformational states which were not apparent for the ACE2 bound RBD, simultaneously reestablishing its dynamic rigidity upon binding. Significantly, the conformational fluctuation between two conformations of the apo-form what we observed can be a reflection of the ‘up’ and ‘down’ conformational changes reported for RBD^4b^.

### Dynamicity based allosteric communication in free RBD

Subsequently, we analyzed the path of free RBD (ACE2-bound RBD was excluded from calculation as it showed negligible cross correlated motions in DCCM) through which allosteric crosstalks propagate from binding loop (selected residue P479) to the identified allosteric loop (selected residue D364). Note that, both the residues were selected based on their highest betweenness-centrality values (Figure S3C) within the highly fluctuating above-mentioned regions. Interestingly, the crosstalk was found to propagate through the hydrogen bonded residues Y423 and D398 in one path (FigureS4A) and P479-D364 *via* A419 to I410 in another path (Figure5A and 3A), all other constituent residues remain the same (see Methods for details). Notably, the constituent residues of the path are evolutionarily conserved^14^. Based on the crystal structure^5^, this hydrogen-bonded communication seems to act as a bridge between binding and allosteric regions of apo-RBD. Hence, we postulate that inherent allostery in apo-RBD is a pivotal factor in ACE2 recognition.

### Hidden electrostatic perturbation revealed favorable allosteric crosstalk in ACE2-bound RBD

The role of hidden contributions (e.g. electrostatic energy perturbation) in molecular recognitions is another important contributor that remains yet to be investigated in case of the spike protein-ACE2 complex. However, structural or dynamic information fail to describe the same and hence, residue wise non-bonded energy contributions in terms of electrostatic 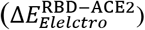 and van der Waals’ 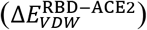 interactions of the RBD towards the ACE2 binding were calculated. Our results successfully captured the binding residue of RBD with significantly high electrostatic energy, particularly, K417. Similarly, binding residues along with some proximal to the binding site showed VDW contributions as expected, for example N487, Q493, Y505, T500, N501 and Y489 (Figure4A-D)^15^. Residue wise electrostatic contributions (up to ~165 kcal/mol) are much higher compared to van der Waals’ (up to ~12 kcal/mol). Hence, undoubtedly in the recognition of ACE2 by RBD, the electrostatic energy is the predominant factor in term of non-bonded energies towards the free energy of binding (Figure S4A). In particular, some significant electrostatic contributions of RBD from distal residues like, D364, K386 etc. were also evident (Figure 4B). We further calculated electrostatic interactions within the RBD in both free and bound states to elucidate the communication path between ACE2 binding residue K417 and allosteric site residue D364 (see Methods for details). Allosteric crosstalk in RBD propagated from K417to D364 *via* R408 to R355 to K356 (Figure 5B). Notably, except binding residue K417,all other constituent partners show evolutionary conservation^14^.

**Figure 4:**
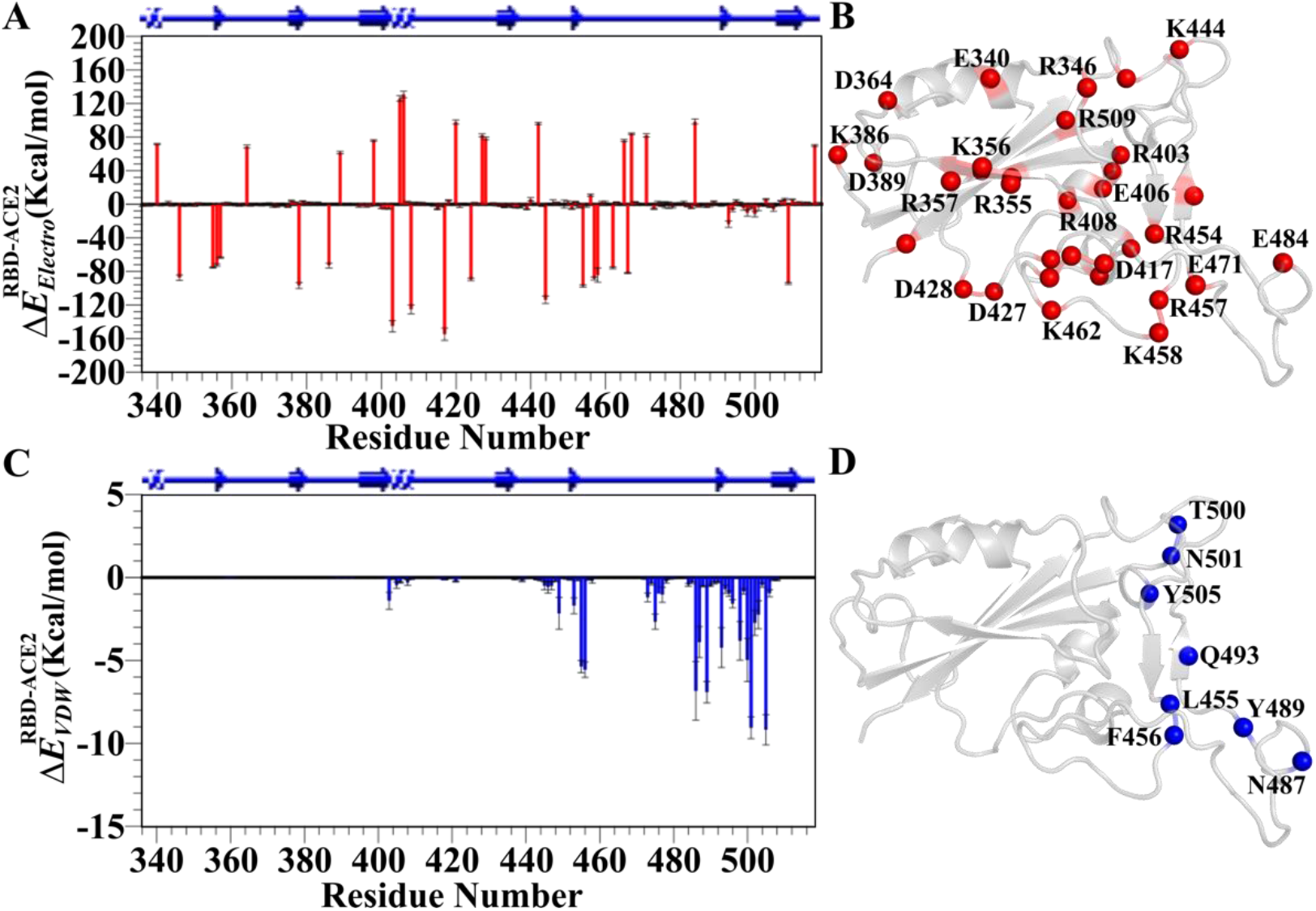
Reside wise nonbonded energy contributions. Residue wise (**A**) electrostatic and (**C**) van der Waals’ contributions of RBD towards ACE2 binding. Contributions of (**B**) electrostatic, and (**D**) VDW were mapped on crystal structures and represented in red and blue spheres respectively.

**Figure 5:**
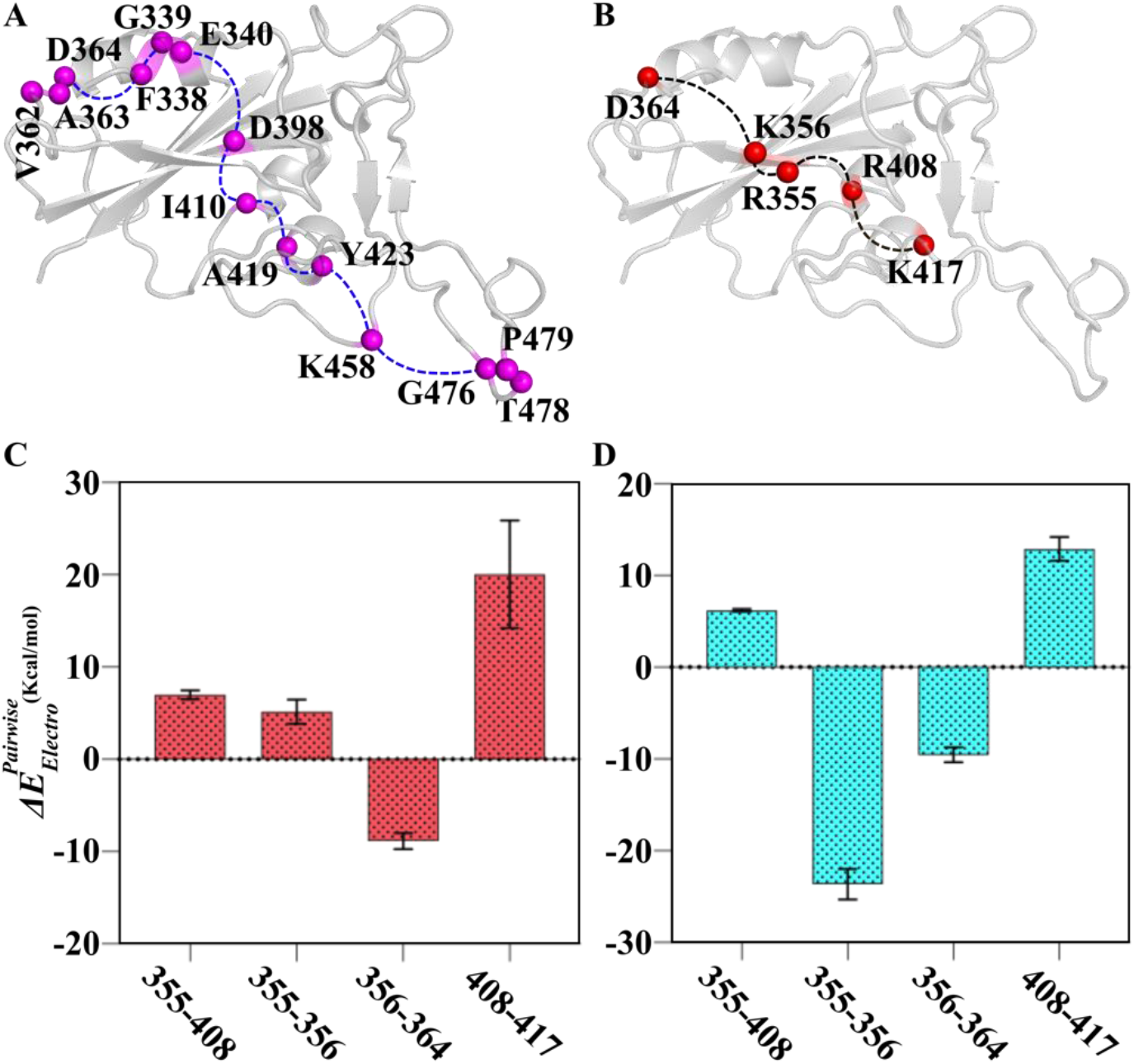
Allosteric communication path and pairwise energy perturbations. (**A**) Path calculated from correlated motions obtained from DCCM. Connecting amino acids on the path (broken blue line) are shown in magenta spheres. (**B**) Represented allosteric crosstalk based on electrostatic energy. Pairwise electrostatic perturbations are shown in free (**C**), and ACE2-bound RBD (**D**). Residues within the path (calculated from electrostatic energy contributions) were selected for pairwise electrostatic energy calculations and represented in red spheres and the path is indicated by broken black line.

Remarkably, the unfavorable pairwise electrostatic interactions of K355-356 pair in apo-form 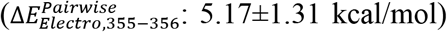 becomes favorable in ACE2-bound form 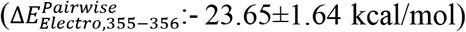. Interestingly, average pairwise contribution for K417-R408 pair of RBD decreased upon complex formation (for free RBD 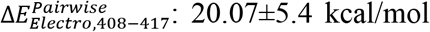 and for ACE2-bound 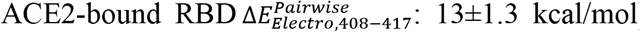) (Figure 5C-D). Conclusively, the results indicated towards the hidden role of pair wise energy perturbation in the recognition of ACE2 by spike protein RBD. Therefore, it can be a site-specific target for allosteric drugs.

Formation of the stable complex can also be verified from the noticeably high residue wise electrostatic contributions compared to the van der Waals’ energy, as reported earlier^15a^. Note that, we mentioned allosteric communication in RBD based on electrostatic energies too, and the pairwise electrostatics become favorable in some pairs after binding to the receptor. On the contrary to dynamic cross-correlation, the existence of favorable communication (between the binding site to the allosteric regions of RBD of one copy towards the S2 domain of another copy) based on pairwise electrostatics point towards hidden transmission of the signal for the cleavage between S1 and S2domain of spike protein following association with ACE2^4a 10.^

### Conclusions

We found that, following binding to ACE2, dynamicity of RBD decreases resulting in stabilization of bound-RBD by the selection of a particular conformation from the fast time scale (ps-ns) regime. While crosstalk mediated by cross-correlated motions between the binding and allosteric loops dominates in free RBD, such dynamic allostery almost disappears when RBD binds to ACE2. In contrast, pair-wise electrostatic energy perturbations dictate favorable allosteric crosstalks within RBD in the ACE2-bound complex. The allosteric paths derived from both dynamic cross correlations and electrostatic energies can be claimed relevant for biological functions considering evolutionary conservation of the residues involved.

The architecture of an individual monomer of spike protein is extended in shape placing S2 domain away from the S1 (comprising RBD). Interestingly, however, the criss-cross arrangement of monomers in trimeric spike places the allosteric site of RBD of one monomer in close proximity of the S2 domain of another copy. Thus, the allosteric communication likely transmits from RBD of one monomer to S2 of another copy in the trimer (Movie S1). It is conceivable that the trimer of spike protein needs to function in a concerted way. Such inter-monomer communication allows regulation of concerted conformational changes within the trimeric spike.

Taken together, our study suggests that the allostery in RBD plays a key role in the process of host cell insertion upon recognition of ACE2 receptor. Based on our results, we propose that allosteric region of the ACE2-binding site of RBD, reported here, can be a potential target for allosteric drug designing. In the context of recent SARS-Cov-2-related outbreak, when designing drug against SARS-Cov-2 is a pressing need, residue-wise dynamics, as well as energy information provided in this study would be beneficial for drug designing purpose.

## Methods

### Molecular Dynamics Simulation studied (MD)

The ACE2-bound (PDB ID: 6M0J, residues 333-518 of RBD and whole ACE2) and free RBD of SARS-Cov-2 crystal structure (extracted from 6M0J, residues 333-518) were equilibrated for 1ns with capped N- and C-termini by N-methyl amide and acetyl groups for both. MD simulations were performed in with both the free and bound spike protein with GROMACS 5.1.2 software^16^ using Amber99SB-Ildn force field^17^ and TIP3P water model^18^ with a box size of 1nm. The titrable residue protonation states were defined using MCCE (multi-conformation continuum electrostatics) method^19^ and also based on PROPKA^20^ web server (http://nbcr-222.ucsd.edu/pdb2pqr_2.0.0) with respect to pH 7.2. The systems were neutralized by adding 150 m MNaCl by substituting appropriate number of solvent molecules to mimic experimental salt condition^15a^. The structures were energy minimized followed by two-step equilibration, namely NVT equilibration followed by NPT equilibration. Temperature was controlled through velocity rescaling^21^ at 310 K with a time constant of 0.1 ps, and pressure was controlled using Parrinello–Rahman barostat^22^ at 1 bar. The particle mesh Ewald algorithm ^23^ was applied to calculate long-range electrostatic interactions. The cut-off for short-range electrostatics and van der Waals’ interactions were 1.2 nm. MD simulations of 1 μs each were performed along with steps were recorded at every 2 ps and analyzed results.

### Dynamic Cross Correlation Mapping (DCCM)

Cross-correlation maps were used to identify the regions that move in or out of phase during the simulations^11^. The elements of the matrix (*C_ij_*) were obtained from their position vector (*r*) as shown in Eq. 1:

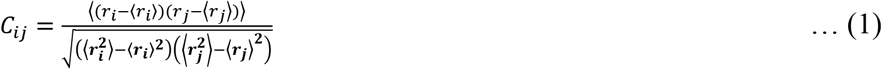

Where *i* and *j* correspond to any two atoms, residues, or domains; *r*_i_ and *r_j_* are position vectors of *i* and *j*; and the angle brackets denote an ensemble average. Inter-atomic cross-correlation fluctuations between any two pairs of atoms (or residues) can be calculated by using this expression and can be represented graphically by the DCCM. The value of *C_ij_* can vary from −1 (strongly anti-correlated motion) to +1 (strongly correlated motion).

### Order parameter (*S*^2^)

Backbone N-H vectors were selected to calculate *S*^2^ over the period of trajectory, which represent dynamicity of protein, with a value 1 indicating complete rigidity and a value towards 0 represents enhanced dynamicity.

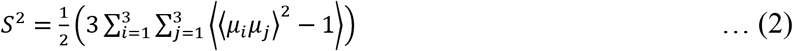

where *μ*_1_, *μ*_2_, and *μ*_3_ are the x, y, and z components of the relevant bond vector scaled to unit magnitude, μ, respectively^9^. Angular brackets indicate averaging over the snapshots.

### Non-bonded Energy calculation

The residue wise non-bonded interaction energy between free RBD and its bound state with ACE2 was described as:

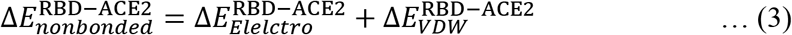

The non-bonded interactions 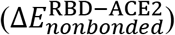 include both electrostatic 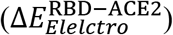 and van der Waals 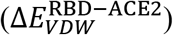 interactions and were modeled using a Coulomb and Lennard-Jones (LJ) potential function, respectively.

### Path analysis from Correlated Motions

Hierarchical clustering was implemented for each correlation network to produce cumulative nodal clusters, or communities those are strongly intra-connectedyet slightly inter-connected, using a related betweenness clustering algorithm to that developed by Girvan.Nevertheless, as is standard for unweighted networks, rather than selecting the partitions with the highest modularity ranking, we selected the partition nearest to the highest modularity score that consisted of the smallest number of overall communities. This excluded the typical circumstance where several small communities with modularity values of similarly high scores were created. The density of connections per node which was assessed by the node centralities were calculated as described by the equation:

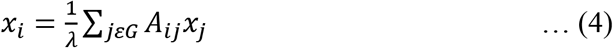

Where *x_i_*, represents the centrality of the node *i*, *A_ij_* is the *ij*th entry of the adjacent matrix *A, λ* is actually a constant, and G represents all nodes. *A_ij_* ≠ 0 if node *i* and *j* are linked, and it will be equal to *e^-w_ij_^*, where *W_ij_* is the edge weight. For every *i* (*i* ∈G) is equivalent to defining the eigenvalues and eigenvectors of matrix A. Considering a pair of nodes optimal (shortest) and suboptimal (near however farther than optimized neo) linking node paths were established employing the previously mentioned algorithm^24^.

### Path analysis from electrostatic energy contributions

The perturbation in pair-wise electrostatic interactions 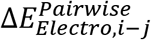:

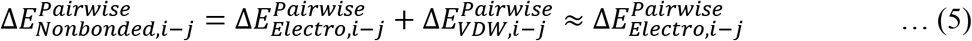

The contributions due to LJ and electrostatic (Coulomb) non-bonded interactions to nonbonded energy,were calculated separately, but the LJ terms were found to be numerically much smaller than the respective electrostatic ones, so we have focused on the electrostatic interactions. The interaction energy between two residues *i* and *j* is the sum of the non-bonded interaction energies already defined in a force-field where:

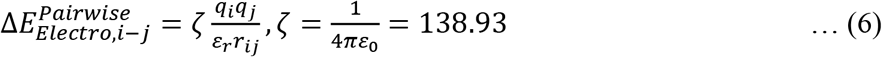

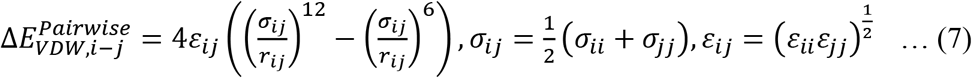

Energy networks were built by considering the amino-acid residues as nodes. A weighted edge was made between any pair of residues *i* and *j* by considering the interaction energy as the weight. Energy-Hubs are defined as nodes that have a higher degree or connectivity in the network.

Betweenness-Centrality was computed using the following equation:

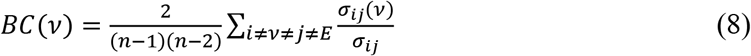

Where *BC(v)* is betweenness-centrality of residue *v.n* is the number of residues within network, *σ_ij_(v)* is the number of shortest paths between residue *i* and *j* that pass through residue *v* and *σ_ij_* is the total number of shortest paths from *i* to *j*. Dijkstra’s algorithm^25^ was applied to calculate the shortest path between residues *i* and *j.*

### Binding free energy

MMPB-SA tool for GROMACS was used to calculate the binding free energy between ACE2 and RBD^26^.

## Author Contributions

SB and JS conceived the project. SB designed and performed all the computational work. SB, RB and JS analyzed the data and wrote the paper.

## Acknowledgments

This work was supported by SERB, DST (India) sponsored project, and CSIR-Indian Institute of Chemical Biology, Kolkata, India. We acknowledge the Central Instrument Facility (CIF) of CSIR-Indian Institute of Chemical Biology, and CSIR-4Pi for super computer facility for supporting our computational work. SB and RB acknowledge UGC and CSIR, India, respectively for awarding senior research fellowship.

## Conflict of Interest Statement

The authors declare no conflict of interest.

## Notes

### Competing Interest Statement

The authors have declared no competing interest.

