## Supporting information for "Transmission of allosteric response within the homotrimer of SARS-CoV-2 spike upon recognition of ACE2 receptor by the receptor-binding domain"

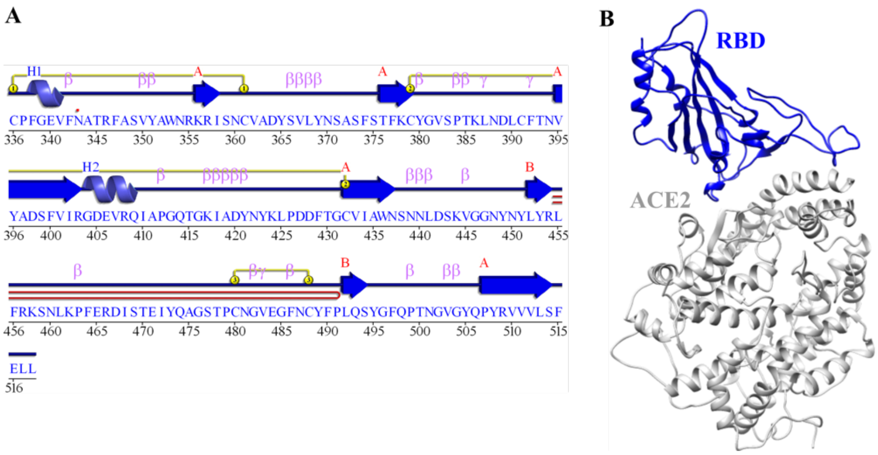


**Figure S1: Structural representation of RBD.** (**A**) Schematic 2D representation of apo-RBD. Disulphide linkages (S-S bonds) are indicated by yellow lines. (**B**) ACE2-bound RBD crystal structure (PDB ID: 6M0J) is shown in cartoon representation.

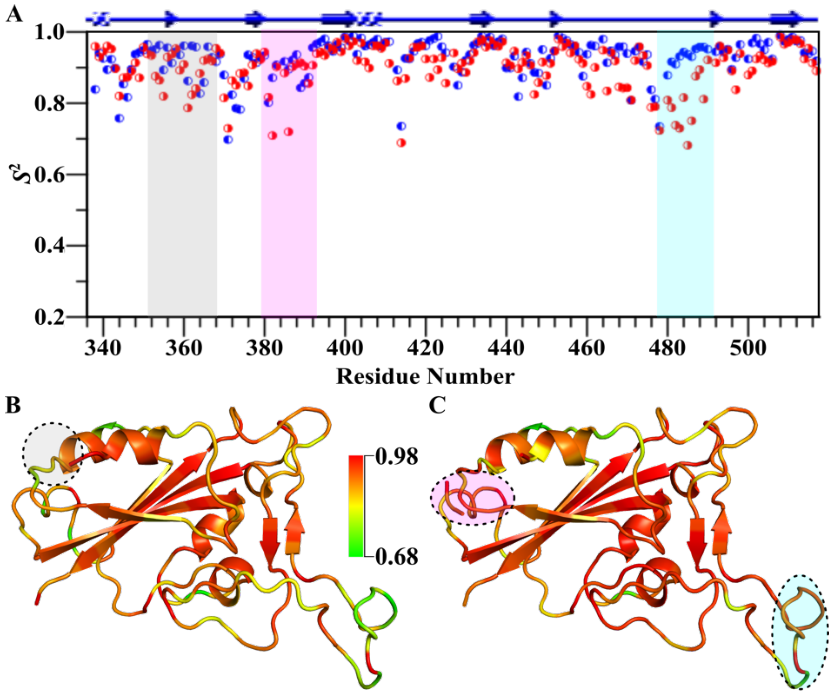


**Figure S2: Order Parameter comparison.** (**A**) $S^2$comparison between apo- (red) and ACE2-bound (blue) RBD. Significant changes are marked with different colored columns. $S^2$ values were mapped onto the crystal structures (PDB ID: 6M0J) of free (**B)**, and ACE2-bound RBD (**C**) respectively. The PyMOL spectrum bar based color scheme was used to point out differential fluctuating regions of both and are indicated by circles correspond to the same color scheme used in (**A**).

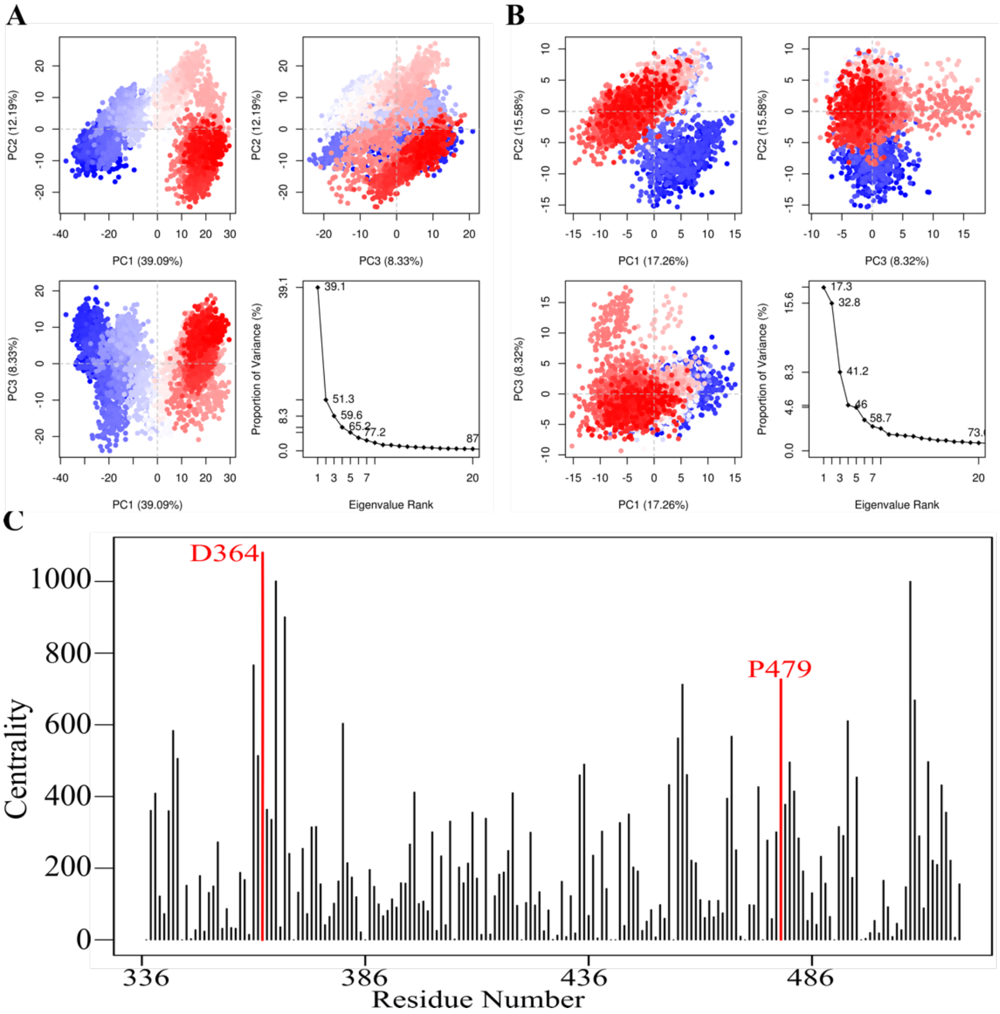


**Figure S3: PCA and Betweenness-centrality of RBD.** PCA projections of different components along with Eigen value rank of apo-form (**A**), and ACE2-bound RBD (**B**). (**C**) Calculated Betweenness-centrality of apo-RBD. Source and sink residues or *vice versa* are marked and indicated in red.

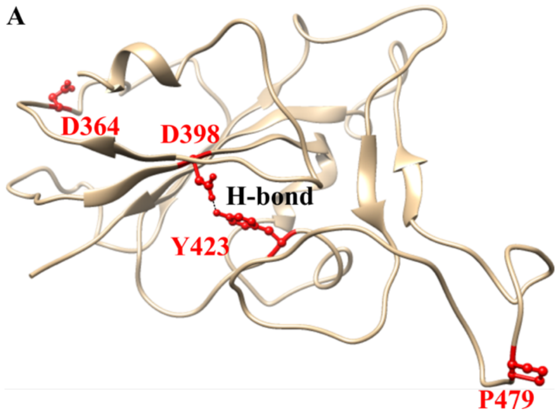


**Figure S4: Correlated residues within path.** (**A**) Represents path captured the distal residue H-bonding which connected the binding region to distal region. Residues were marked in ball-stick model as well as colored and labeled in red. Hydrogen bond is show in black broken line. Other residues of the path (shown in Figure 5) are not shown here.

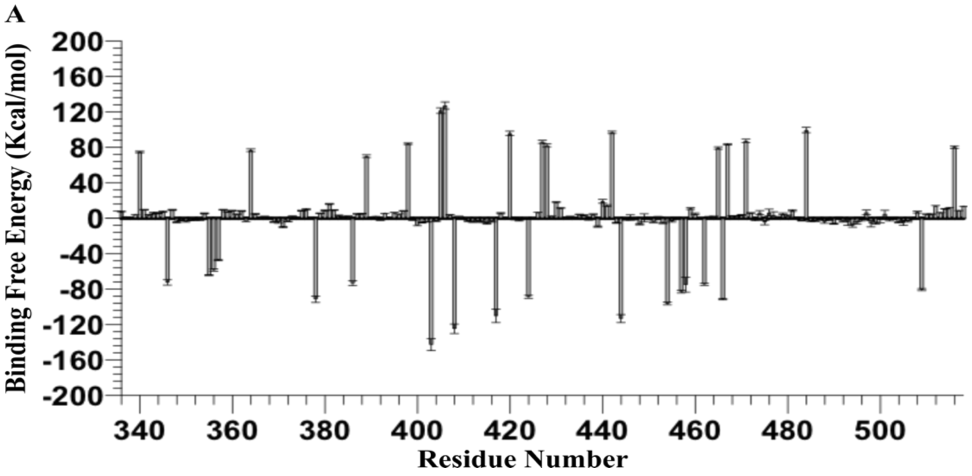


**Figure S5: Binding free energy.** (**A**) Residue wise free energy of binding of ACE2-bound RBD.

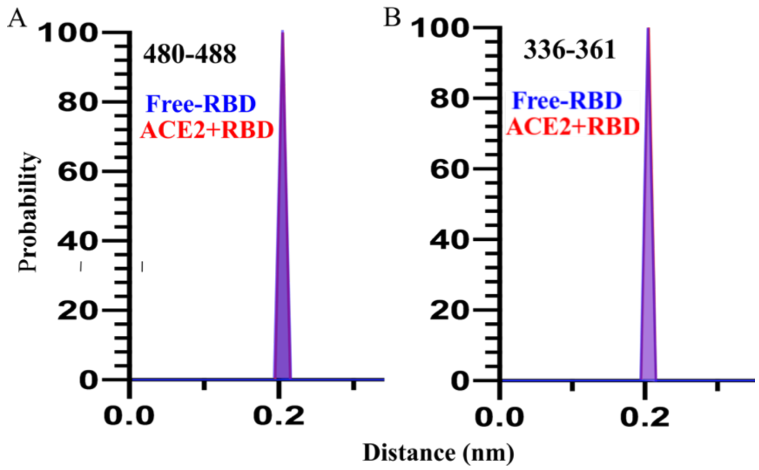


**Figure S6. Disulfide bond population**. Population at 0.2 nm clearly indicated the retention of the S-S bond throughout the simulation in (A) C480-C488, and (B) C336-C361 pairs. Note that these S-S bonds are evolutionarily conserved.

**Legend for the Supporting Movie S1:**

Conformational alteration of allosteric loop along with binding loop derived from final simulated structures (i.e.1μs state) of both apo- and ACE2-bound RBD. Corresponding residues e.g. D364 from allosteric loop and K417 from binding loop are labelled and marked in red ball stick. RBD (cartoon representation in magenta) from one monomer are in contact with S2 domain of another monomer (shown in blue cartoon form, representative residue D985 shown by blue ball and stick), when seen in trimer constitution, through another distal loop (corresponding residue N388, marked in red ball and stick), indicating the allosteric post-binding (with ACE2, represented in green cartoon) signal transfer for subsequent processes.

**Softwares used**

Following softwares were used to create images for all the illustrations.

INKSCAPE 0.92 (https://inkscape.org/release/0.92.2/mac-os-x/), UCSF Chimera[26],

PyMOL (The PyMOL Molecular Graphics System, Version 2.0 Schrödinger, LLC.),

GraphPad Prism8 (https://www.graphpad.com/scientific-software/prism/),

GIMP(https://www.gimp.org/downloads/), Bio3d[27], MMPBSA[25],

Servers:PDBsum

(http://www.ebi.ac.uk/thornton-srv/databases/cgi-bin/pdbsum/GetPage.pl?pdbcode=index.html)
